# *Exiguobacterium* Sp. HA2, isolated from the Ilam Mountains of Iran

**DOI:** 10.1101/2021.03.12.435112

**Authors:** Reza Heidari, Mostafa Akbariqomi, Gholamreza Tavoosidana, Garshasb Rigi

## Abstract

A motile, Gram-stain-positive, rod-shaped, non-sporing, tolerate up to 5% NaCl, grew at 0–25 °C, designated *Exiguobacterium* sp. HA2 was isolated from the soil of the Ilam Mountains of Iran during October 2016. The major isoprenoid quinone is MK-7 and in the smaller amount are MK-6 and MK-8. Polar lipids included diphosphatidylglycerol, phosphatidylglycerol, phosphatidylserine, phosphatidylinositol, phosphatidylethanolamine. Major fatty acids (>10 %) are isoC_13:0_, isoC_15:0_ and C_16:0_. The bacterial cell wall peptidoglycan layer was lysine-glycine. The 16S rRNA sequence was analyzed at the phylogenetic levels. Also, A supplemental comparison was made between five other genes including *csp, gyrB, hsp70, rpoB,* and *citC.* According to the results of genotypic and phenotypic characteristics, the strain was categorized in the genus *Exiguobacterium.* This bacterium had the closest relation with *Exiguobacterium undae,* and thus was dubbed *Exiguobacterium* sp. HA2. The different in the Phenotypic, functional characteristics and phylogenetic indicated *Exiguobacterium* sp. HA2 can be regarded as representing considered a novel species within the genus *Exiguobacterium*.

## Introduction

The genus of *Exiguobacterium* includes psychrophilic, thermophilic and mesophilic strains and species [1]. The *Exiguobacterium* genus is anaerobic, gram-positive bacteria of low G+C contents. This bacterium has been repeatedly isolated from the ancient Siberian sediments, as well as environments such as Greenland glacial, the spa springs (Yellowstone National Park), and food processing environments. Therefore, the isolation sources of this bacterium are totally different and often in extreme environments [2]. Accordingly, each of these species or strains appears to have specifically and exclusively adapted to the environments. The diversity in their genome has been responsible for their adaptation to different conditions [3].

This genus was divided into two groups according to the 16S rDNA sequence and genomic sequence; group I is the psychrophilic *Exiguobacterium,* and group II thermophilic [4]. Many strains of *Exiguobacterium* have unique properties that make them suitable for applications in biotechnology, detergent industry, sewage treatment, and agriculture [5]. Psychrophilic microorganisms have been described that are capable of growing at <20°C [6].

In the present study, *Exiguobacterium* was isolated from the soil samples collected from the Ilam Mountains (one of Iran’s cold provinces), and its genotypic and phenotypic characteristics were assessed and identified.

## Isolation, cultivation conditions and maintenance of strain

*Exiguobacterium* sp. HA2 has been isolated from soil samples by cold enrichment method. The soil samples were collected from the top layer of soil (5 cm) from an area in Ilam (33° 45’ 30.32’’ N, 46° 11’ 51.33’’ E). The soil sample was store at cold room at 4 °C for 72h before isolation. For isolation, 10 g of soil sample was added in 50 ml sterile water, and then the suspension was incubated at 4°C for 60 min. The strain was isolated on Tryptone Soy Agar (TSA; Difco Laboratories, Detroit, MI, USA) at 4°C for 5 days. The isolated bacterium was stored at −80°C in TSB (Tryptone Soya Broth) supplemented with 25% glycerol for cryoprotectection.

## Genomic characterization

Genomic DNA was extraction and purification using the THP(Triton/Heat/Phenol) slight modification of protocol [7]. Also, Phylogenetic comparisons were performed between the mentioned new strain and sequences of five other genes including *csp* (universal major cold shock protein), *gyrB* (gyrase subunit B), *hsp70* (Class I-heat shock protein-chaperonin), *rpoB* (DNA-directed RNA polymerase beta subunit), and *cit*C (isocitrate dehydrogenase). To confirm bacterial identification, we amplified the genes (16s rRNA, *csp*, *gyrB, hsp70, rpoB, citC)* directly by using the universal primers for *Exiguobacterium* **(Table1)**. The annealing temperatures were respectively 56, 53, for *csp*, *rpo*B, and 49°C for *citC* and *gyr*B, and 47°C for *hsp70.* The PCR amplified products were purified using QIAquick Kit (PCR Purification Kit, Qiagen, Valencia, CA, USA) and then sequenced [8]. The TOPO TA Cloning Kit (Life Technologies) was used for the cloning and sequencing of the amplicons of other genes. The Qiagen Plasmid Kit (Qiagen, Valencia, CA, USA) was then used to extract the obtained clones. An ABI 373a DNA sequencer was used for the analysis of the products of cycle sequencing performed by a Perkin-Elmer 9600 thermal cycler and an ABI Dye Terminator Chemistry (PE Applied Biosystems).

Genotypic analyses were conducted to clarify the phylogenetic relationship between the new strain and the most relevant set of five reference strains. The sequence analysis of the 16S rRNA gene indicated that new strain was categorized in the genus *Exiguobacterium* **(Fig. 1)**. The sequence was deposited in GenBank with the accession number KT967971. The sequences of *gyr*B, *citC, rpoB, hsp70,* and *csp*, also were deposited in GenBank with the accession numbers of KX574228, MH370480, MH378445, MH378775, and MH379139, respectively. Phylogeny analysis of other genes was performed to confirm identification the phylogenetic relationships of this strain within *Exiguobacterium.* Phylogenetic trees constructed based on *gyr*B **(Fig. 2)**, *citC* **(Fig. 3)**, *rpo*B **(Fig. 4)**, *hsp70* **(Fig. 5)**, and *csp* **(Fig. 6)** sequences of the following genes. Analysis of the 16S rRNA gene sequences indicates that HA2 had a high level of similarity to SH3 and undae (96%). Also, the alignment and comparison sequence of *gyr*B, *citC, rpoB, hsp70, csp* of HA2 showed highest similarity to *Exiguobacterium* undae. Analysis of the gene sequences of HA2 with other members of the genus *Exiguobacterium* indicated that similarities with phylogenetic neighbours were in the range 60-100%.

**Fig. 1.**
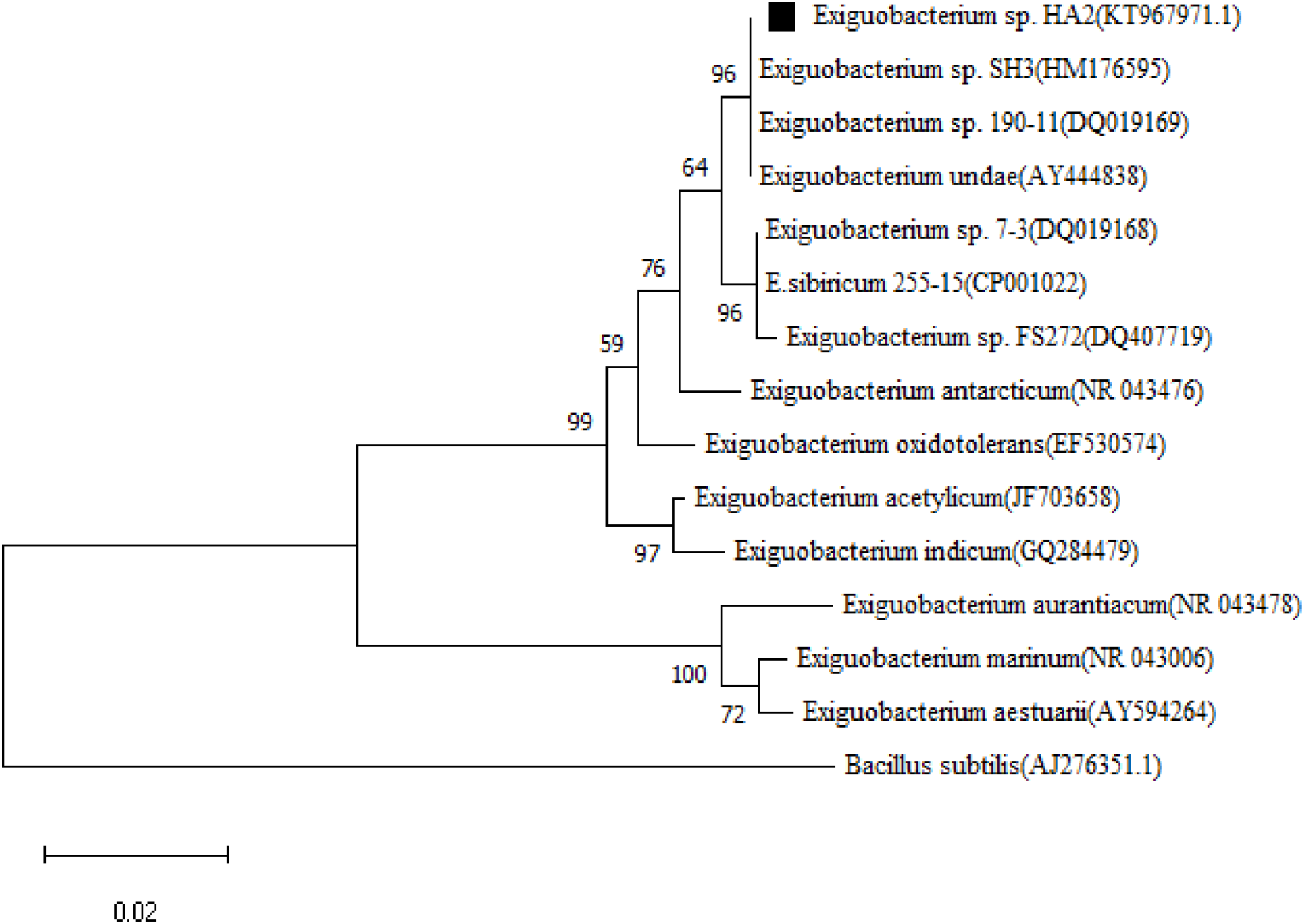
Phylogenetic tree based on 16S rRNA gene sequence. Numbers at the nodes indicate the bootstrap values on neighbour joining analysis

**Fig. 2.**
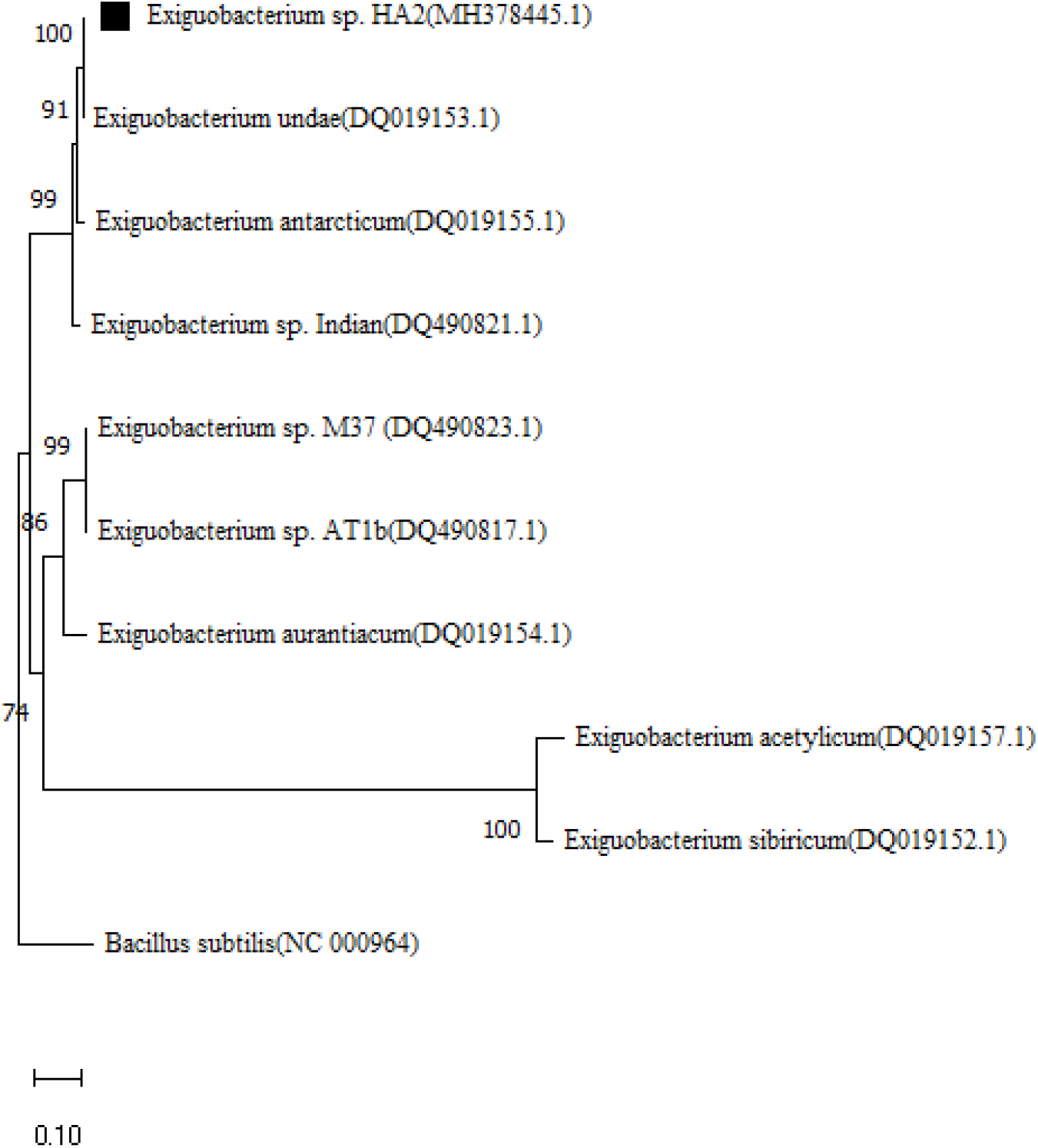
Phylogenetic tree based on *gyr*B gene sequence. Numbers at the nodes indicate the bootstrap values on neighbour joining analysis

**Fig. 3.**
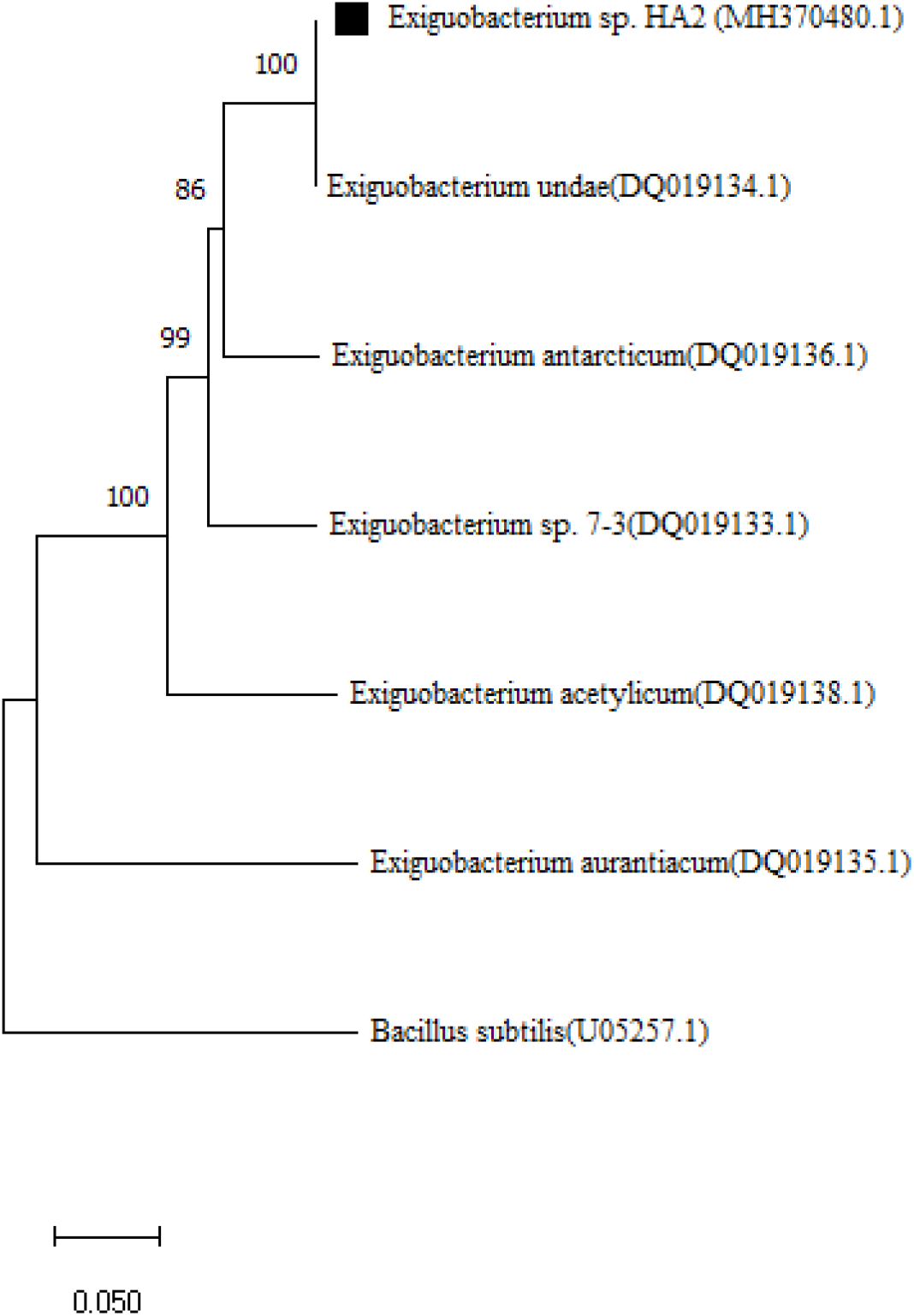
Phylogenetic tree based on *citC* gene sequence. Numbers at the nodes indicate the bootstrap values on neighbour joining analysis

**Fig. 4.**
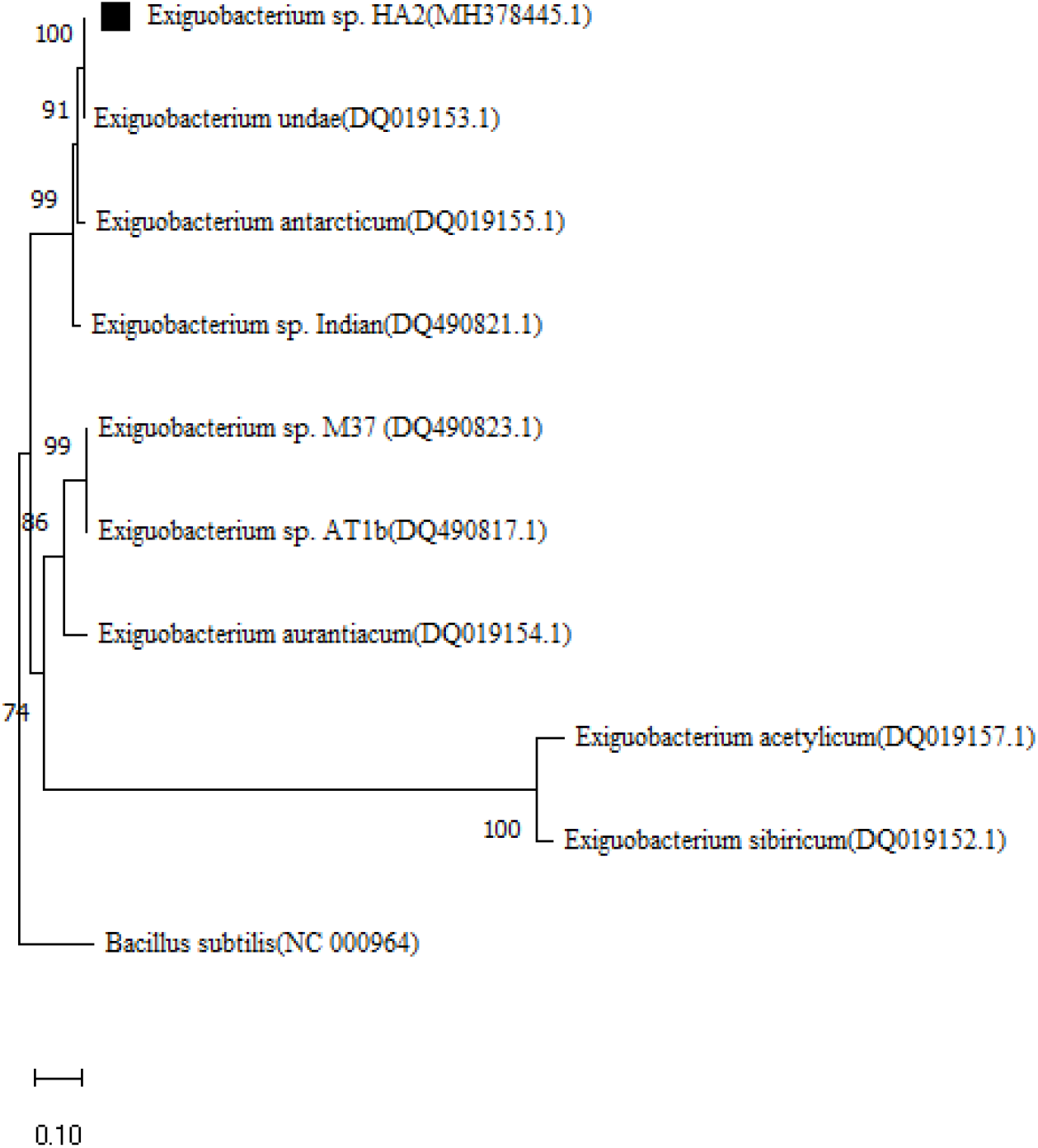
Phylogenetic tree based on *rpoB* gene sequence. Numbers at the nodes indicate the bootstrap values on neighbour joining analysis

**Fig. 5.**
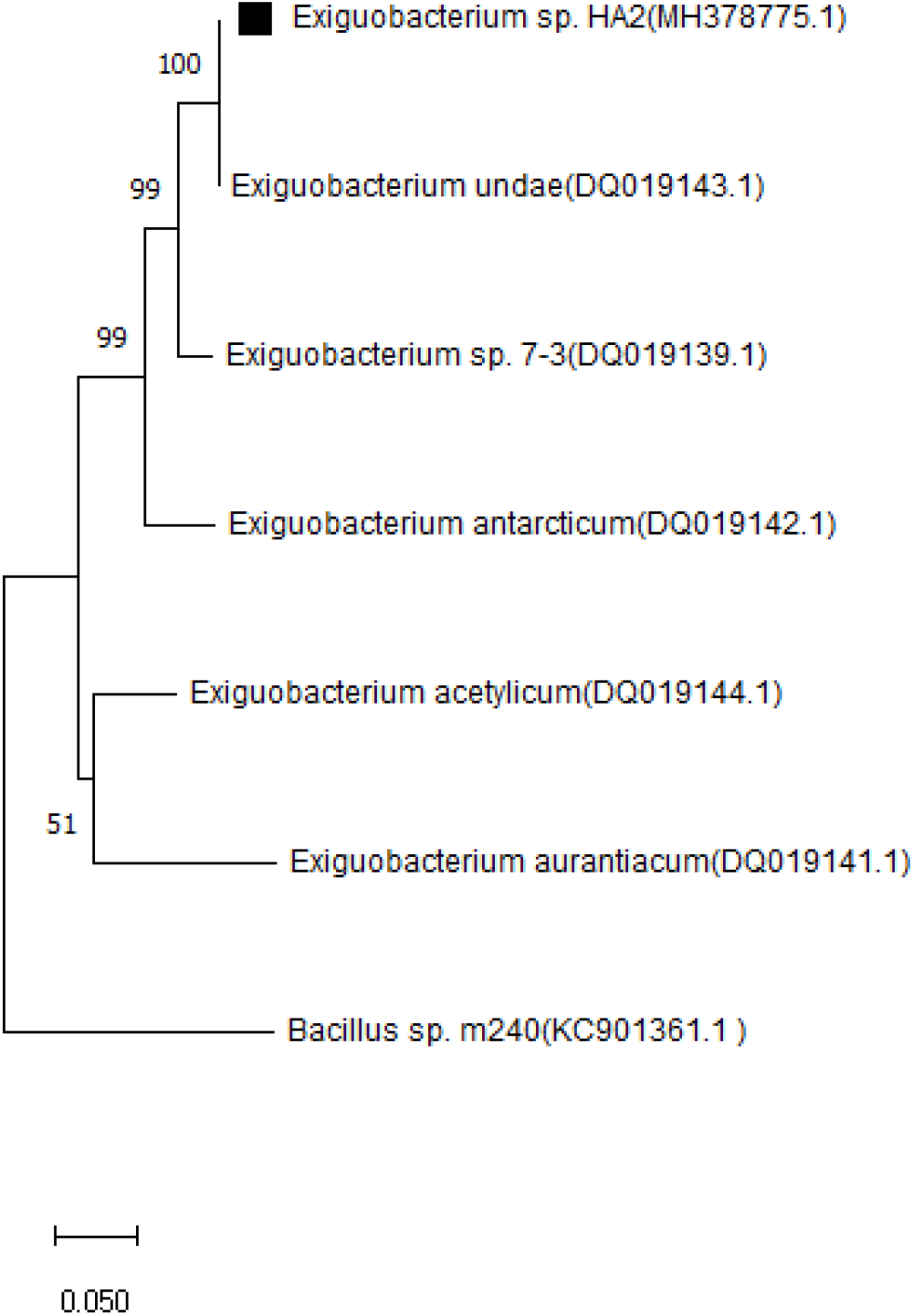
Phylogenetic tree based on *hsp70* gene sequence. Numbers at the nodes indicate the bootstrap values on neighbour joining analysis

**Fig. 6.**
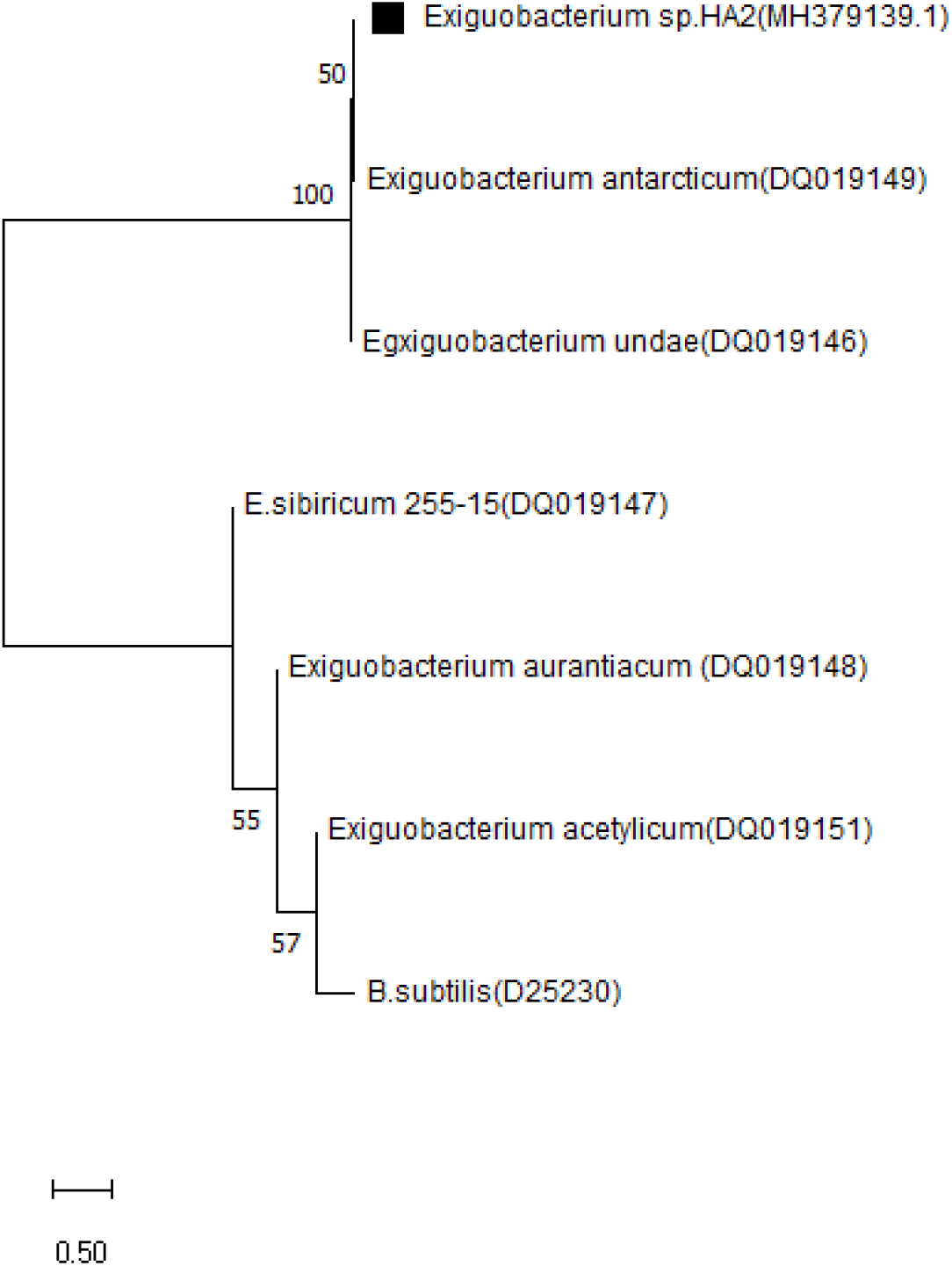
Phylogenetic tree based on *csp* gene sequence. Numbers at the nodes indicate the bootstrap values on neighbour joining analysis

Using the MEGA5.0 program, we constructed Phylogenetic trees based on the neighbor-joining and distance methods. The phylogenetic tree was reconstructed with generated by the maximumlikelihood method and the reliability of each node was assessed with 1000 bootstrap test replications. The scale bar represents 0.1 substitution per nucleotide position. The roots of the trees were determined using those genes from *Bacillus*.

## Morphology, physiology and biochemical characterization

The optimum temperature required for the growth of *Exiguobacterium* sp. HA2 was 15°C. A colony size desirable for scanning electron microscopy (SEM) was reached by incubating inoculated R2A plates at 4°C for 28 days. Samples for scanning electron microscopy (SEM) studies were prepared according to the previously published method [9]. Gram-staining was performed as described previously [10]. Minimum and maximum temperatures for growth was investigated on R2A agar after incubation for 5 days at −4 to 30°C (−4, 0, 4, 10, 15, 20, 25, 30°C). Tolerance to NaCl was determined by incubating R2A broth supplemented with NaCl (0–10%, w/v, at 0.5% intervals) concentrations in 1.5 ml. The pH range was evaluated using R2A broth containing various pH 6–10 (at intervals of 0.5 pH unit) prior to autoclaving using appropriate buffers [11]. Presence of spores was evaluated by staining with malachite green. API 20NE, API ID 32GN and API ZYM (bioMerieux) were used for analyzing the production of different enzymes. Fatty acid extraction and analysis was determined by gas chromatography (GC) as described previously [12]. Antibiotic sensitivity testing was performed using standard methods [13]. The culture’s ability to assimilate different carbon compounds was assessed using minimal medium [K2HPO4 2%(w/v); KH2PO4 0.5% (w/v); agar 1% (w/v)]. The analysis of chemotaxonomic markers was performed through cell wall amino acids [14], polar lipids [15], peptidoglycan structure [16] and isoprenoid quinones [17]. *E. sibiricum* DSM 17290, *E. antarcticum* DSM 14480T, *E. oxidotolerans* JCM 12280T, *E. acetylicum* DSM 20416T, *E. undae* DSM 14481T, and were used as reference strains in morphological and biochemical assessments and fatty acids identification.

Total the membrane lipoquinones and polar lipids were extracted and separated according to the previous described [18] phospholipids were visualized by staining with molybdenum blue spray (Sigma). The peptidoglycan was isolated and analyzed using a method adapted from Schleifer and Kandler [14]. after hydrolysis the cells, total-cell sugars were analysis by TLC on cellulose plates according to the previous methods [19]. The major isoprenoid quinone is MK-7 and in the smaller amount are MK-6 and MK-8. Polar lipids compositions including diphosphatidylglycerol, phosphatidylglycerol, phosphatidylserine, phosphatidylinositol, phosphatidylethanolamine. Major fatty acids (>10 %) are iC13:0, iC15:0 and C16:0; minor components are listed in **Table 2**. The peptidoglycan type is lysine-glycine.

**Table 1.**
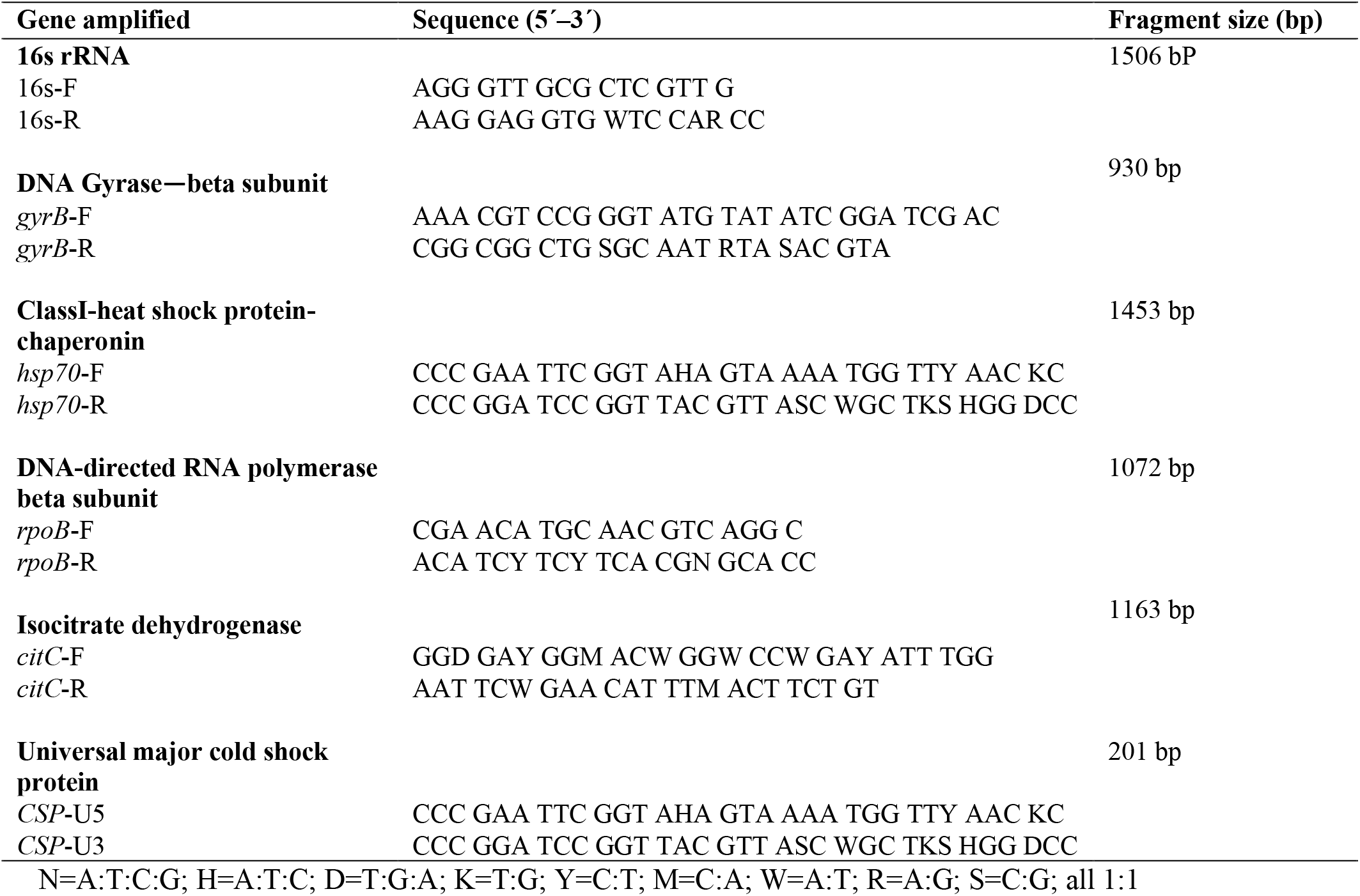
Primers used for PCR amplification for *Exiguobacterium* genus analysis.

**Table 2.** Fatty acid composition of the Siberian permafrost isolates and the type strains of Exiguobacterium

|  | HA2 | 7-3<sup>b</sup> | 255-15<sup>b</sup> | 190-11<sup>b</sup> | *E. undae*<sup>a</sup> | *E. antarcticum*<sup>a</sup> | *E. aurantiacum*<sup>a</sup> |
| --- | --- | --- | --- | --- | --- | --- | --- |
| iC11:0 |  |  |  |  |  |  | 2 |
| iC12:0 | 3 | 2 | 2 | 2 | 2 | 3 | 3 |
| C12:0 | 1 | 1 |  | 1 |  | 1 | 2 |
| iC13:0 | 11 | 9 | 13 | 8 | 9 | 12 | 18 |
| AiC13:0 | 8 | 11 | 15 | 10 | 9 | 11 | 12 |
| iC14:0 | 9 | 1 | 1 | 1 | 2 | 1 |  |
| C14:0 | 2 | 3 | 1 | 2 | 2 | 2 | 3 |
| iC15:0 | 11 | 13 | 12 | 13 | 10 | 11 | 4 |
| aiC15:0 | 4 | 3 | 4 | 3 | 3 | 2 |  |
| C16:1–11c | 9 | 8 | 3 | 7 | 8 | 18 | 10 |
| iC16:0 | 3 | 2 | 2 | 2 | 2 |  |  |
| C16:1–5c |  | 1 |  | 1 |  |  |  |
| C16:1–7c |  | 1 |  | 1 | 7 | 3 |  |
| C16:0 | 16 | 17 | 20 | 12 | 17 | 13 | 27 |
| C17:1–10c | 4 | 2 | 2 | 3 | 2 | 3 |  |
| iC17:0 | 5 | 9 | 12 | 8 | 7 | 5 | 6 |
| aiC17:0 | 3 | 3 | 3 | 3 | 2 |  |  |
| C18:1–9c |  | 2 | 1 | 2 | 3 | 6 | 2 |
| C18:1–7c | 2 | 2 | 1 | 3 | 3 |  |  |
| C18:0 | 5 | 4 | 5 | 3 | 6 | 5 | 5 |

Strains: 1, *Exiguobacterium* sp. HA2; 2, *Exiguobacterium* strains 7-33; 3, *Exiguobacterium* strains 255-15; 4, *Exiguobacterium* sp. 190-11; 5, *E. undae* DSM 14481T; 6, *E. antarcticum* DSM 14480T; 7, *E. aurantiacum* DSM 6208T.
Only values >1% are indicated; values ≥5% are given in bold
aData obtained from Rodrigues et al. (2006) [20]
bData obtained from Fruhling et al. (2002) [21]

Cells of strain HA2 were Gram-positive, non-sporing, motile, facultatively anaerobic, rod-shaped, approximately 0.8–1μm in width and 1.5–2μm in length **(Fig. 7)**. Growth occurred at between 0 and 25°C (optimally, 15°C), pH grows at pH 6-10 (optimally, 7.0) and in the presence of 0-12 (w/v) NaCl with an optimum of approximately 3 (w/v) NaCl. positive in tests for oxidase, Gelatinase, b-galactosidase, DNase, catalase, Caseinase, Phosphatase, Lysine decarboxylase, but are negative for, urease and H2S production and for the indole test. *Exiguobacterium* sp. HA2 utilize D-Fructose, D-Galactose, D-Mannose, L-Rhamnose, Cellobiose, D-Lactose, Maltose, Starch, Amygdalin, Arbutin, Glycogen, Citrate utilization. Strain HA2 was susceptible to Amikacin (30μg), Amoxicillin (30μg), Clindamycin (25μg), Colistin (10μg), Doxycycline (25μg), Co-trimoxazole (25μg), Nalidixic acid (30μg), Norfloxacin (10μg), Nitrofurantoin (300μg), Sulfamethoxazole (50μg), tobramycin (15μg), lomefloxacin (30μg), roxithromycin (30μg), ciprofloxacin (30μg), lincomycin (15μg), cefotaxime (30μg), cefazolin (30μg), kanamycin (30μg), novobiocin (30μg), chloramphenicol (30 μg), ampicillin (25μg), tetracycline (30μg), streptomycin (25μg), erythromycin (15μg), bacitracin (10μg), gentamicin G(30μg), polymyxin B (50 μg), oleandomycin (15μg), spectinomycin (100μg), rifampicin (25μg) and carbenicillin (100μg). but resistant to ceftriaxone (30μg) norfloxacin (10μg), gentamycin (10μg). The phenotypic features and biochemical profile of HA2 compare with other reference strains are description and illustrated in **Table 3**.

**Fig. 7.**
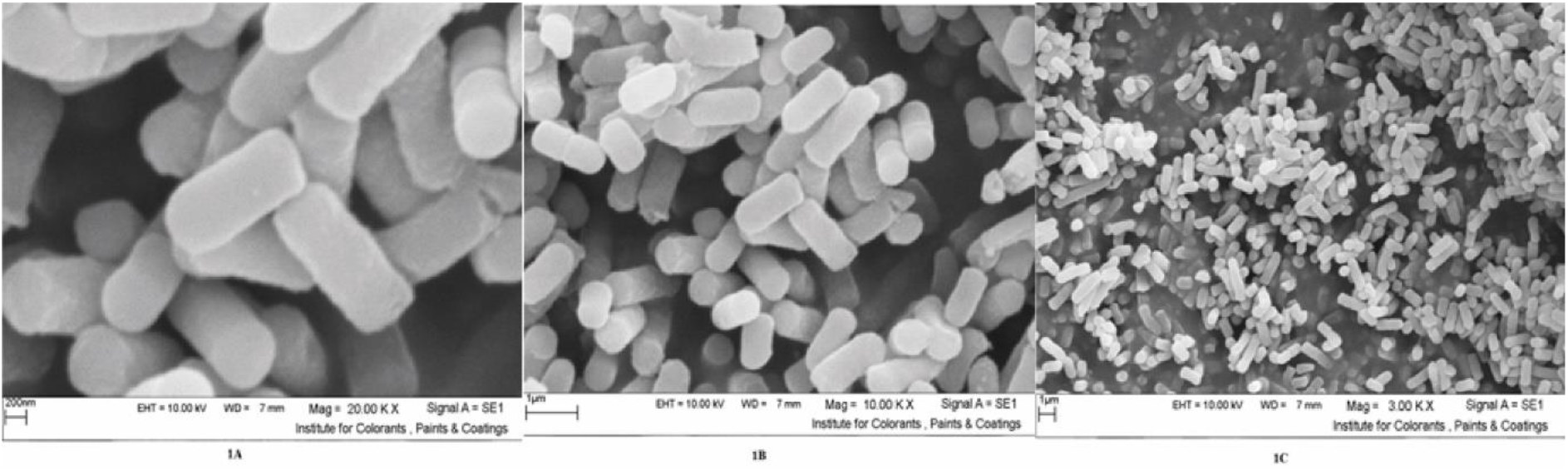
Scanning electron microscopy images of *Exiguobacterium* sp. HA2 with different resolutions

**Table 3.**
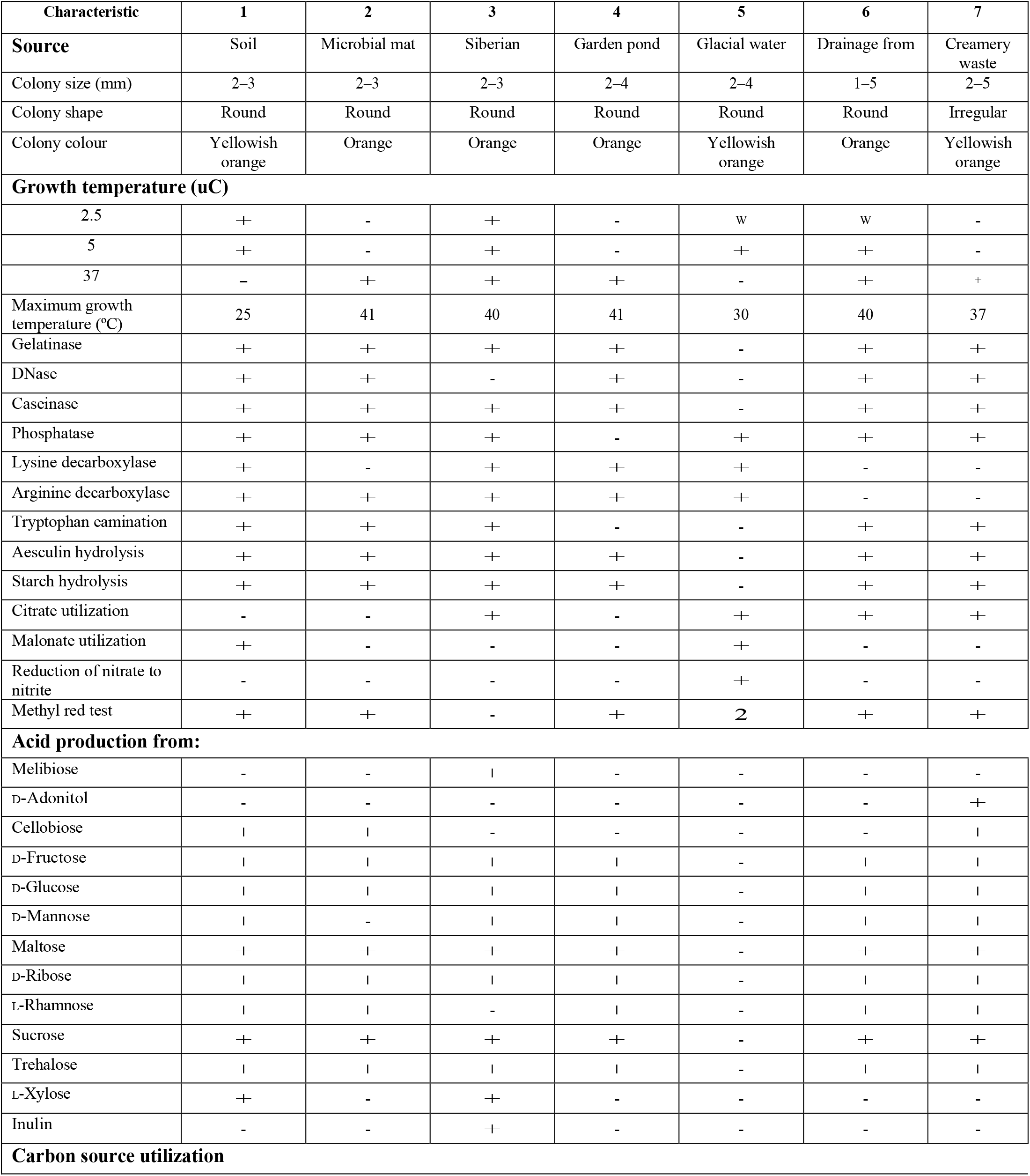
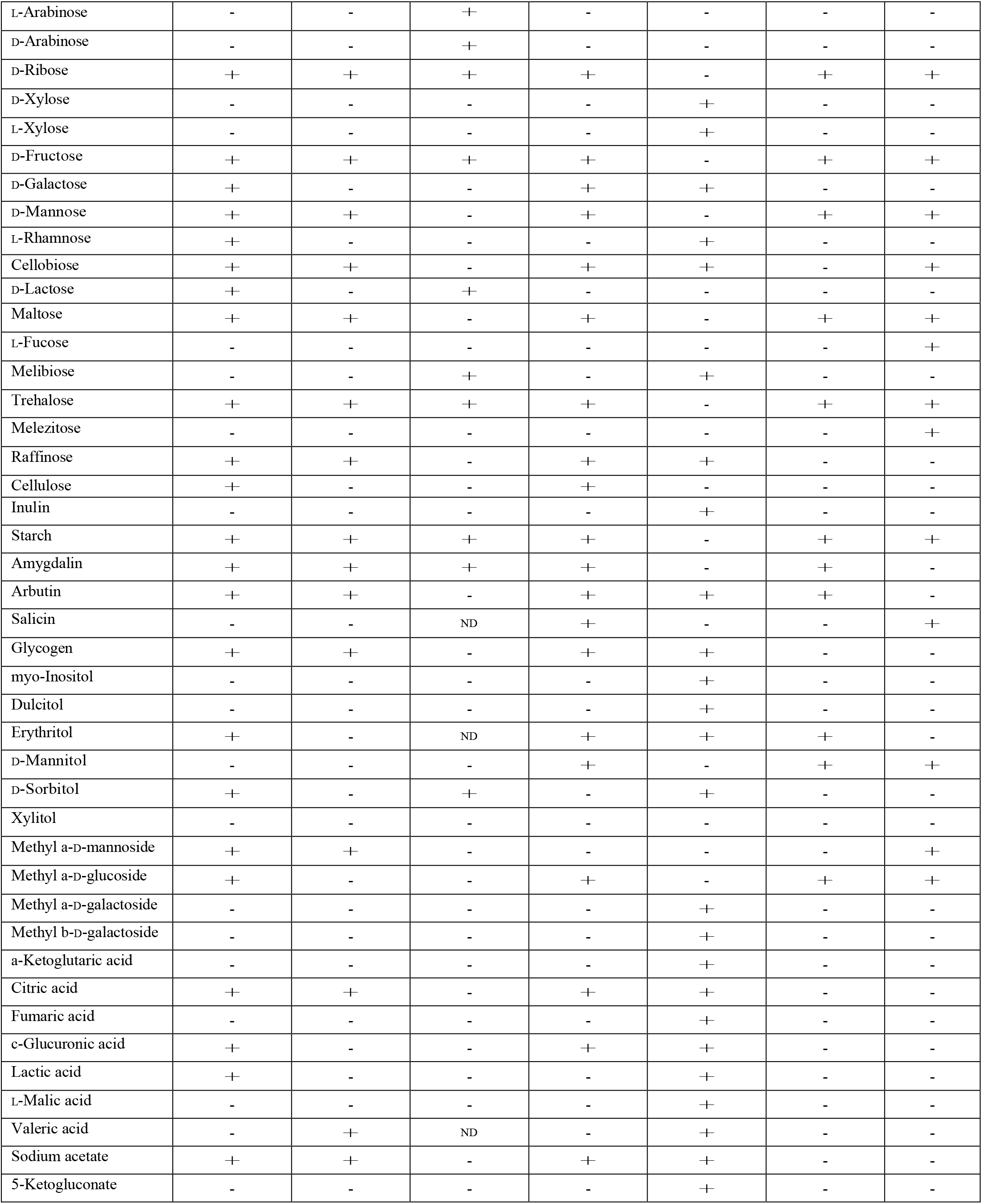
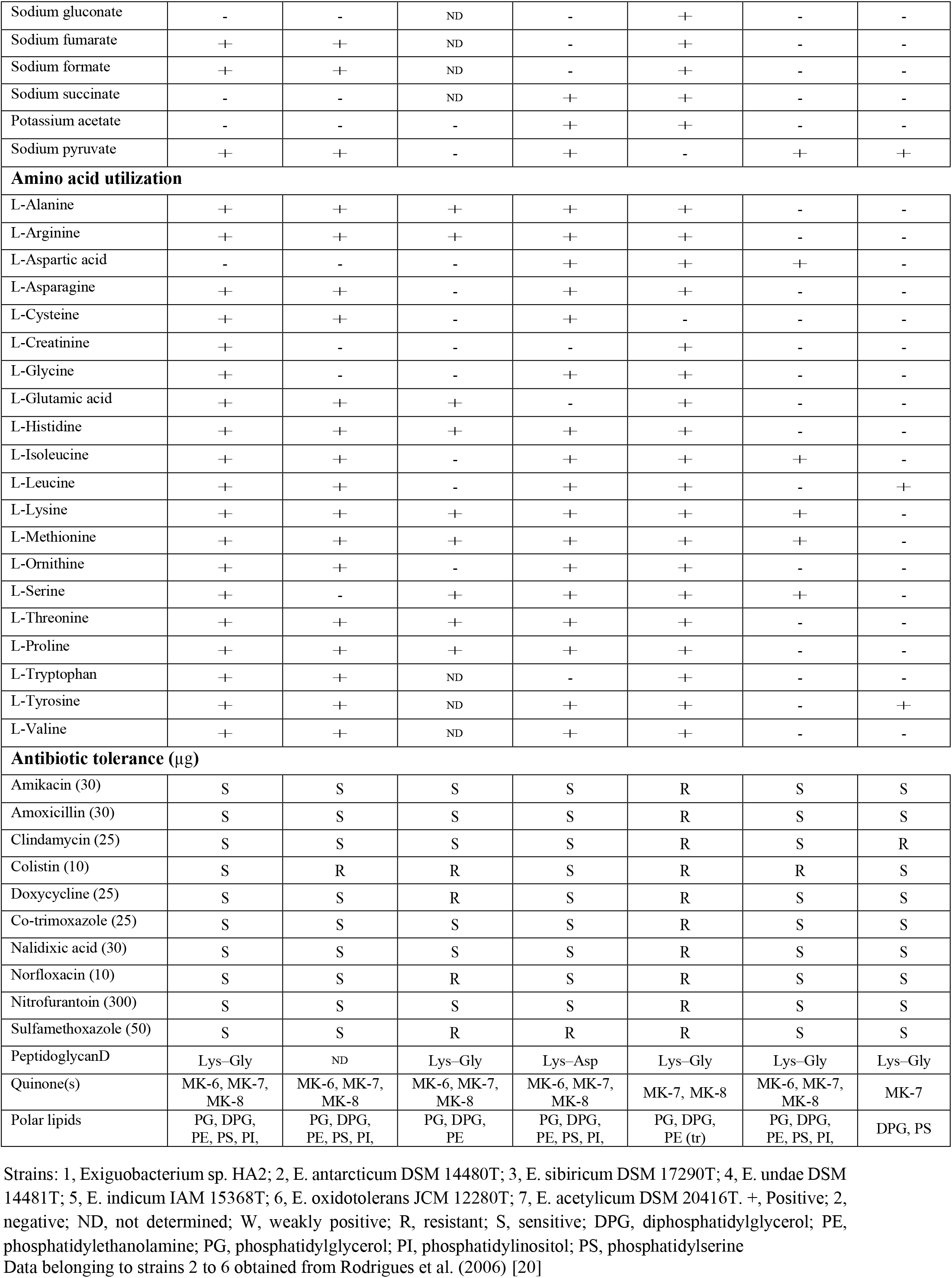
Phenotypic features distinguishing strain *Exiguobacterium* sp. HA2 from the six most closely related species of the genus *Exiguobacterium*

## Description of *Exiguobacterium* sp. HA2

Phenotypic evaluations were also conducted to clarify the phenotypic distinguishability of the new isolates from each other and the reference *Exiguobacterium* strains. Scanning electron microscope images of the strain are shown in Fig. 7. Cells are Gram-positive, rod-shaped (2–3mm), aerobic, non-spore-forming. As presented in Table 1, surface colonies are bright, yellowish orange, convex, entire and shiny. Growth occurs at 0 to 25°C; optimal temperature for growth about 15°C. Tolerate to 5 % NaCl, grow at pH 6–10, are positive in tests for oxidase, Gelatinase, b-galactosidase DNase, catalase, Caseinase, Phosphatase, amylase, lipase and protease. *Exiguobacterium*. sp. HA2 utilize D- glucose, sucrose, starch, Trehalose, Raffinose, Cellulose, Amygdalin, Arbutin, Glycogen, D- Sorbitol, Citric acid, Lactic acid. Cells are sensitive to the following antibiotics (μg): Amikacin (30) Amoxicillin (30) Clindamycin (25) Colistin (10) Doxycycline (25) Co-trimoxazole (25) Nalidixic acid (30) Norfloxacin (10) Nitrofurantoin (300) Sulfamethoxazole (50). The peptidoglycan type is lysine-glycine. The major isoprenoid quinone is MK-7 and in the smaller amount are MK-6 and MK-8. Polar lipids included diphosphatidylglycerol, phosphatidylglycerol, phosphatidylserine, phosphatidylinositol, phosphatidylethanolamine. Major fatty acids (>10 %) are iC13:0, iC15:0 and C16:0; minor components are listed in Table 2. Isolated from a soil from Ilam, Iran.

## Conflicts of interest

The authors declare that there are no conflicts of interest.

## Ethical statement

This article does not contain any studies with animals performed by any of the authors.

## Notes

### Competing Interest Statement

The authors have declared no competing interest.

